# Functional Network Connectivity Imprint In Febrile Seizures

**DOI:** 10.1101/2021.12.18.473280

**Authors:** Ullas V Acharya, Karthik Kulanthaivelu, Rajanikant Panda, Jitender Saini, Arun K Gupta, Bindu Parayil Sankaran, Kenchaiah Raghavendra, Ravindranath Chowdary Mundlamuri, Sanjib Sinha, ML Keshavamurthy, Rose Dawn Bharath

**Author notes:** Equal contributing authors (Ullas V Acharya, Karthik Kulanthaivelu, Rajanikant Panda). **Corresponding author** Dr Rose Dawn Bharath, D.M., Professor and Head, Department of Neuroimaging and Interventional Radiology, Mailing address: Department of Neuroimaging and Interventional Radiology, National Institute of Mental Health and Neurosciences, Bengaluru-560029, Karnataka, India.

## Abstract

Complex febrile seizures (CFS), a subset of paediatric febrile seizures (FS), have been studied for their prognosis, epileptogenic potential and neurocognitive outcome. We evaluated their functional connectivity differences with simple febrile seizures (SFS) in children with recent-onset FS. Resting-state fMRI (rs-fMRI) datasets of 24 children with recently diagnosed FS (SFS-n=11; CFS-n=13) were analysed. Functional connectivity (FC) was estimated using time series correlation of seed region–to-whole-brain-voxels and network topology was assessed using graph theory measures. Regional connectivity differences were correlated with clinical characteristics (FDR corrected *p <* 0.05). CFS patients demonstrated increased FC of the bilateral middle temporal pole (MTP), and bilateral thalami when compared to SFS. Network topology study revealed increased clustering coefficient and decreased participation coefficient in basal ganglia and thalamus suggesting an inefficient-unbalanced network topology in patients with CFS. The number of seizure recurrences negatively correlated with the integration of Left Thalamus (r= −0.58) and FC measures of Left MTP to ‘Right Supplementary Motor and left Precentral’ connectivity (r=−0.53). The FC of Right MTP to Left Amygdala, Putamen, Parahippocampal, and Orbital Frontal Cortex (r=0.61) and FC of Left Thalamus to left Putamen, Pallidum, Caudate, Thalamus Hippocampus and Insula (r 0.55) showed a positive correlation to the duration of the longest seizure. The findings of the current study report altered connectivity in children with CFS proportional to the seizure recurrence and duration. Regardless of the causal/consequential nature, such observations demonstrate the imprint of these disease-defining variables of febrile seizures on the developing brain.

## INTRODUCTION

Febrile seizures (FS), are defined as seizures in children between 6 months to 5 years of age, accompanied by fever, but without evidence of underlying central nervous system (CNS) infection ^1^. The definition excludes seizures in children due to neuroinfection, prior afebrile seizure, or a pre-existent CNS abnormality ^2^. It is the most common type of seizure in childhood, occurring in 2–5% of children ^3^. A polygenic basis for febrile seizures is well elaborated with multiple candidate genes /loci described ^4,5^. Febrile seizures can be subtyped as either Simple Febrile Seizures (SFS) or Complex Febrile Seizures (CFS). SFS, accounting for two-thirds of FS, is typified as a generalized convulsive episode of seizure without features suggesting a focal nature. They typically last for lesser than 15 minutes and do not recur within 24 hours. CFS usually demonstrate any of the following features: focal features; prolonged duration (characterised as lasting more than 15 minutes), seizure recurrence within 24 hours/index febrile illness; postictal neurological deficits ^6,7^.

The immature brain is different from the adult brain in its vulnerability to seizures. It is attributable, in part, to overexpression of factors enhancing neuronal excitability thereby resulting in a relative imbalance of excitation vs inhibition. This phenomenon is critical to activity-directed synaptogenesis during development ^8^. A pertinent concern then is whether FS in a growing brain can influence the neurocognitive outcome. A propos SFS, prospective epidemiological studies have demonstrated a benign outcome, without cognitive dysfunction or subsequent epilepsy^9,10^. Most children who experience SFS do not develop structural mesial temporal abnormalities ^11^. Epilepsy frequency following SFS is estimated at 1.0-2.2 %, a figure not any different from the normal population^12^. The relationship of CFS with temporal lobe epilepsy (TLE), however, is ridden with conflicting data. Retrospective risk-factor analyses have revealed that patients with intractable TLE happen to report a history of prolonged febrile seizures (CFS) at higher frequency (30-60%) ^9,13^. CFS is associated with a heightened risk of epilepsy in 4.1-6.0 % of the cases ^12^. Min Lan Tsai et al, in a study on long term neurocognitive outcomes in subjects with CFS, noted significantly lower full-scale intelligence quotient (FSIQ), perceptual reasoning index, and working memory index scores than in the control group^14^. Dube et al indicated that hyperthermic seizures in the immature rat model of FS do not cause spontaneous limbic seizures during adulthood. ^15^. However, “prolonged” experimental FS led to later-onset limbic (temporal lobe) epilepsy and interictal epileptiform EEG abnormalities in a significant proportion of rats^16^.

The literature on the clinical profile of FS notwithstanding, imaging literature in FS is sparse. Theodore et al., in a study of 35 subjects presenting as refractory Complex Partial Seizures [CPS] with video-EEG of temporal lobe onset found 9 patients with a prior presentation of CFS had smaller volumes of ipsilateral Hippocampal Formation [HF]. FS in the clinical history had a predictive value on the severity of HF atrophy ^17^. Suffices to say that the imaging literature in FS has at best summarised the morphological aspects of the brain, with little emphasis on the functional connectivity, especially in the case of CFS, which seems to have a different evolution. Imaging studies attempting to understand the connectivity differences even in patients with generalised epilepsy are riddled with complexities as the observed differences between patients and matched controls could be secondary to various disease defining features like the type of seizures, duration of disease, number of recurrences, familial loading of epilepsy etc. Hence there is an increased interest in studying drug naïve patients with new-onset seizures and the use of disease matched controls in an attempt to reduce the complexity of the question to be answered^18,19^.

With a theoretical background of dissonant profiles of SFS and CFS from an experimental, clinical and neurocognitive perspective we identify a knowledge gap about early brain connectivity changes in children with CFS vis-à-vis SFS. We hypothesize that there could be differences in the connectivity patterns between clinical subgroups. It is likely that the alteration in connectivity may not be limited to a particular brain region, but may manifest as whole-brain connectivity changes. Graph theory is a relevant formalism in this context for studying large-scale brain networks. Here networks are conceptually represented as sets of nodes (vertices exemplified by ROIs in the brain) and connected by links (edges illustrated by structural, functional or effective connections)^20^. We use seed-region to whole-brain-voxel time-series functional connectivity and Graph theoretical analysis on Blood Oxygen Level Dependent (BOLD) resting-state functional magnetic resonance imaging (rs-fMRI) to compare subjects with CFS and SFS.

## RESULTS

24 subjects were included in the final analysis [Simple FS - 11 and Complex FS - 13] within 12 days [IQR= 8.5 - 13.5] and 10 days [IQR = 9-30] after the last seizure respectively. Among the clinical variables analysed, namely mean age at onset, the number of recurrent febrile seizures, the maximum duration of febrile seizures, duration of the disease, none showed significant between-group differences. Functional connectivity (FC) was estimated using time series correlation of seed region–to-whole-brain-voxels and the brain regions showing significant between-group connectivity differences were correlated with disease-defining clinical characteristics. Our results revealed that the CFS group had altered temporal lobe connectivity and altered basal ganglionic integration measures proportional to recurrences, and duration of the seizure.

### Altered brain connectivity in complex febrile seizure

Seed to Voxel functional connectivity analysis revealed that the patterns of connectivity of multiple seed ROI’s in patients in the CFS group were significantly different (FDR-corrected, P < 0.05 for 90 ROIs) from those in the SFS group (Fig 1, Table 2, Supplementary Figure 1 (a-k)). Patients with CFS demonstrated increased connectivity involving bilateral middle temporal pole (MTP) with left subcortical structures (putamen, amygdala, Accumbens), parahippocampal gyrus, left insula and frontal orbital cortex when compared to the SFS group. (Table 2). Bilateral thalami as seed regions showed increased bilateral connectivity with the subcortical areas [putamen (0.000001), caudate (0.000001), pallidum(0.000001), thalamus (right 0,00258; left 0.000001)], bilateral insular, frontal opercula cortex (Table 2). On the other hand, decreased functional connectivity of the right MTP with the left parietal/central opercular cortex (0,000002, 0,000002) and left post-central gyrus (0,000002) was noted. The left MTP also showed decreased FC with the left precentral (0,002824), right supplementary motor cortex(0,002824) and left central opercular cortex (0,003058).

**Figure 1.**
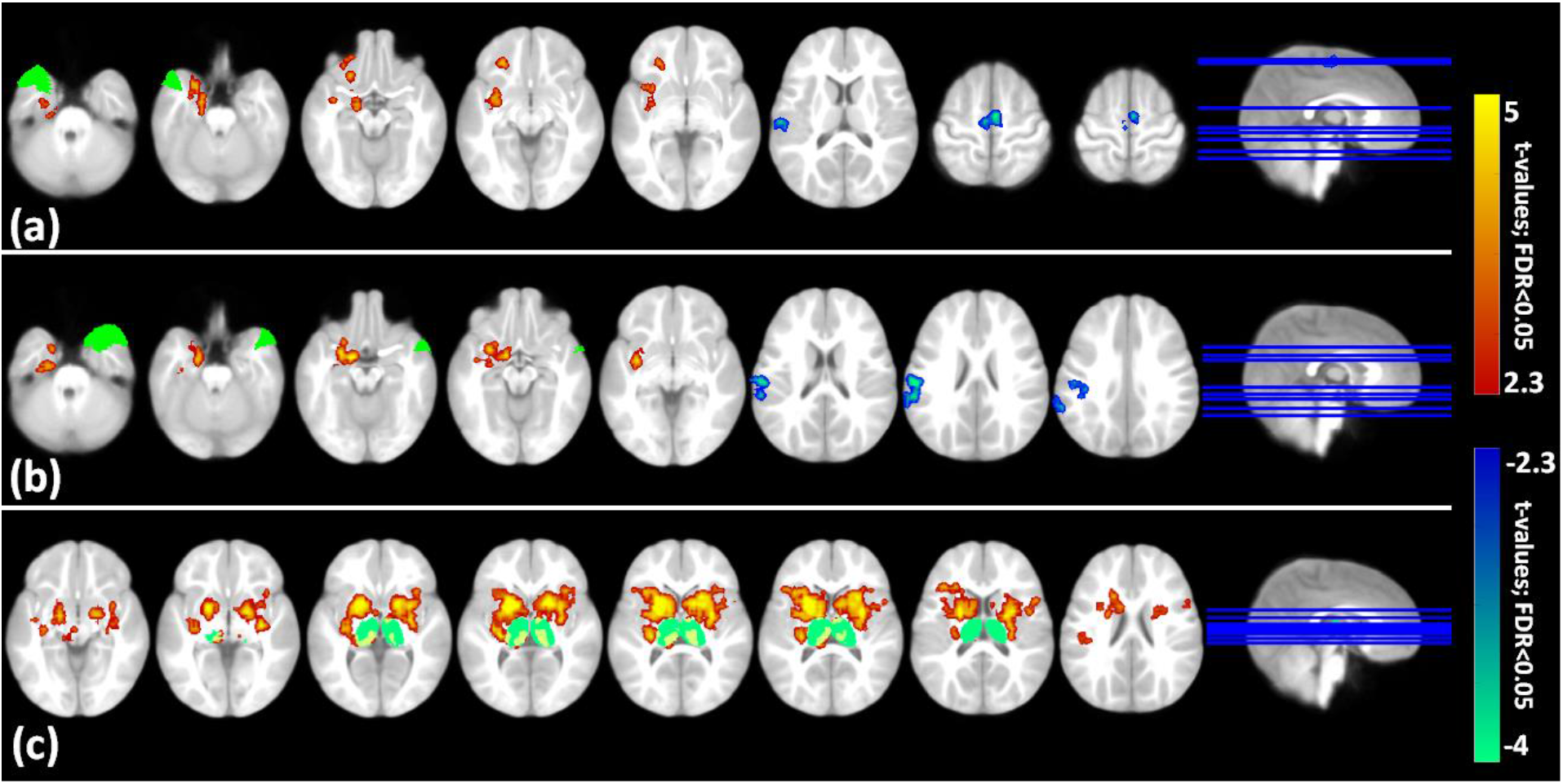
Multi-slice view of significant (FDR corrected; p <0.0.5) seed to voxel-based functional connectivity for Complex (CFS) vs Simple febrile seizure (SFS) of (a) left middle temporal pole (b) right middle temporal pole (c) bilateral thalamus. Red to yellow colour indicates increased connectivity and blue to green colour indicates decreased connectivity in CSF in comparison with SFS. The colour bar indicates “FC t-values” of the connectivity voxels and the dark green cluster in the connectivity map shows the seed region.

### The network topology in complex febrile seizure

The CFS group revealed increased network segregation (ie, clustering coefficient), decreased network integration (i.e., participation coefficient), and decreased global efficiency (Fig 2, Table 3, Supplementary Figure 1 (l-u)). A significantly (FDR-corrected, P < 0.05 for 90 ROIs) increased network segregation was observed in right inferior frontal gyrus (p=0.0005) and left caudate(p=0.001). Bilateral caudate (right hemisphere p=0.0005, left hemisphere p= 0.0032), pallidum (p=0.0009), right thalamus (p=0.0023), Amygdala (p=0.0016), orbitofrontal gyrus (p=0.0004) and paracentral lobule (p=0.0027) revealed decreased network integration (Fig3; Table 3).

**Figure 2.**
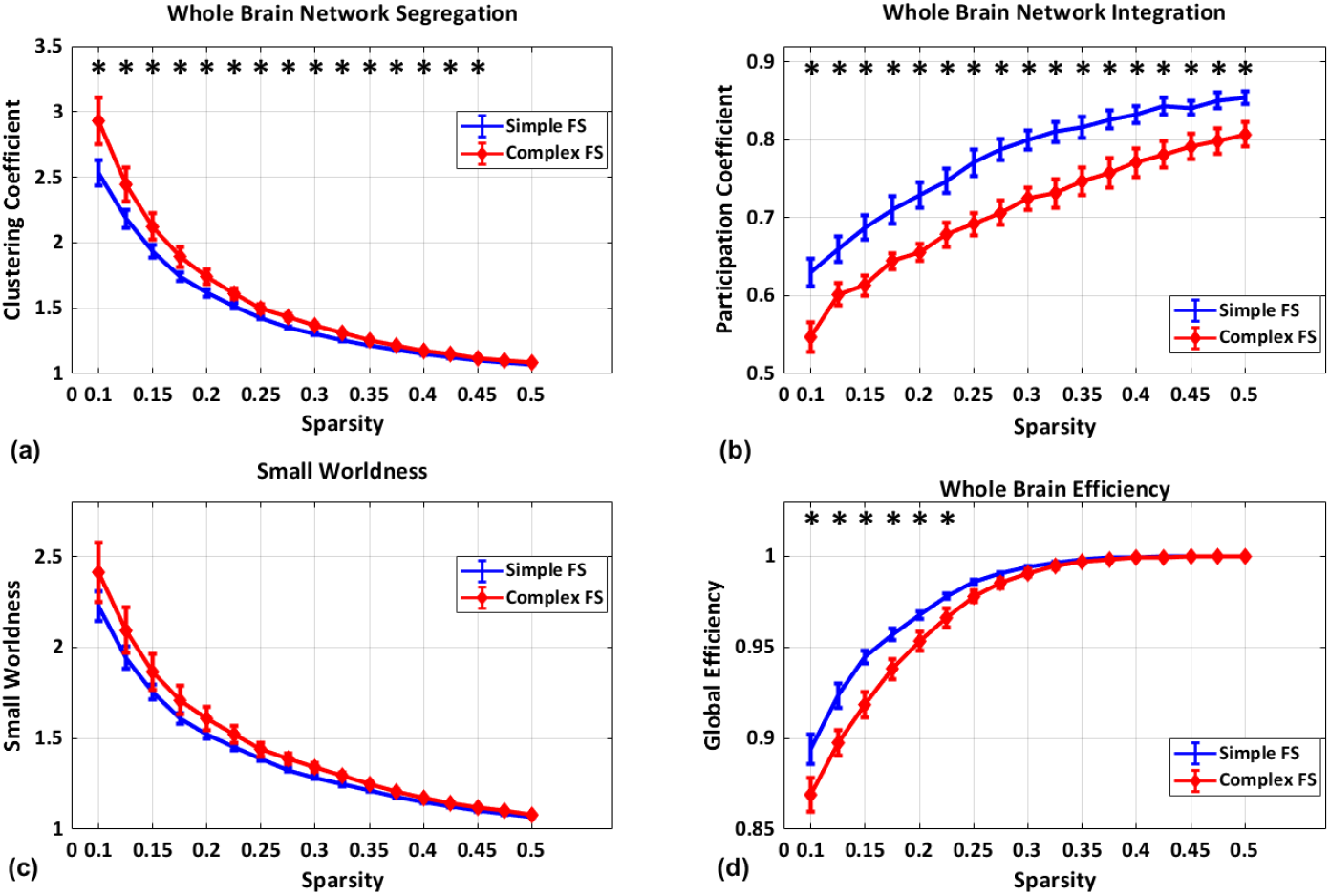
Whole Brain network topological properties. average across all ROIs as determined by Graph theoretical analysis for (a) network segregation (b) network integration (c) small worldness (d) global efficiency for complex febrile seizure (CFS) (red line graph) and simple febrile seizure (blue line graph). The error bar in the graph represents a 95% confidence interval and the star represents a significant difference between complex and simple febrile seizure with correction for multiple comparisons; FDR<0.05 for the number of sparsity (N=17).

**Figure 3.**
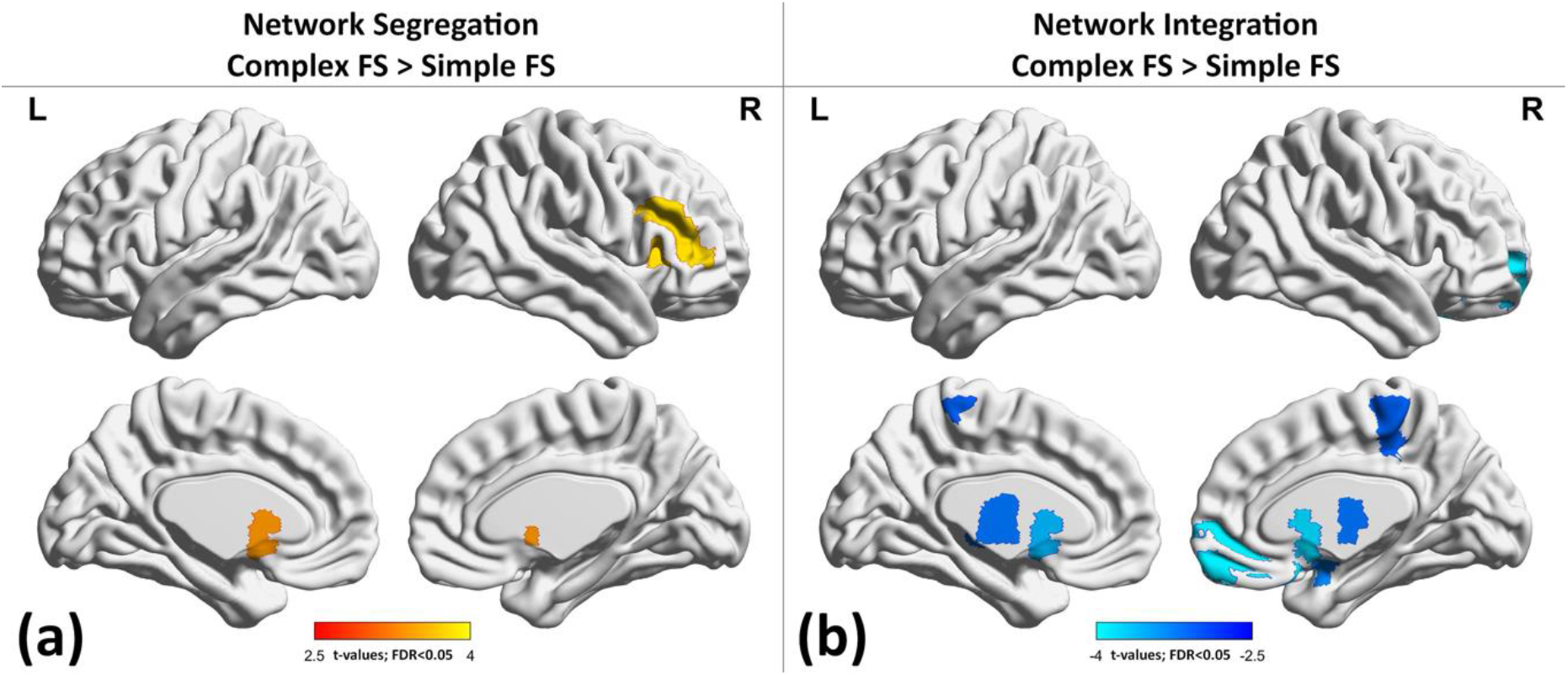
Glass brain view of group difference between CSF and SFS for regional network segregation and integration. (a) brain regions showed higher segregation/local connectivity (i.e., nodal clustering coefficient) in CFS compared to SFS and (b) brain regions showed decreased integration (i.e., nodal participation coefficient) in CFS compared to SFS with multiple comparisons correction of FDR<0.05 for no of ROIs (N=90). Red to yellow color indicates increased network segregation and blue to green color indicates decreased network integration in CSF in comparison with SFS. The color bar indicates the “t-values” of statistical difference between CFS and SFS. The glass brain view were constructed using BrainNet Viewer.

### Correlation with clinical variables

The number of recurrences of febrile seizures was negatively correlated with connectivity between Left MTP to Right Supplementary Motor and left Precentral Cluster (r= −.53 and p =0.011) (Figure 4C). A similar finding was observed with measures of integration of the left thalamus (as derived by Graph theoretical analysis) (r= −0.58 and p =0.004) (Figure 4D).

**Figure 4.**
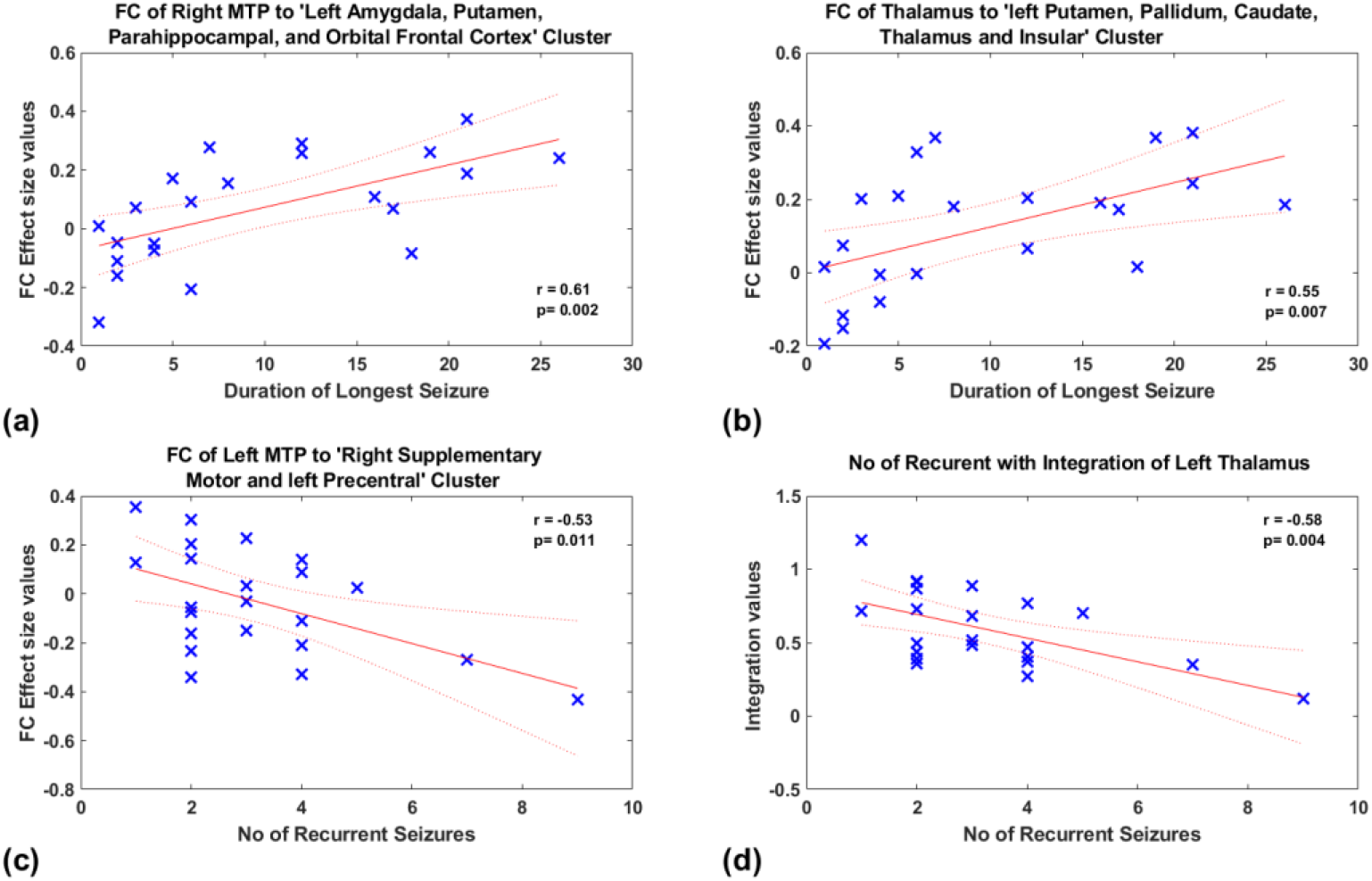
Correlation of FC and Network topology with clinical variables of all CFS and SFS subjects. The FC effect size is the mean t-values of every voxel in the corresponding cluster.

Furthermore, the duration of the longest febrile seizures was positively correlated (Figure 4A) with the functional connectivity of Right MTP to Left Amygdala, Putamen, Parahippocampal, and Orbital Frontal Cortex Cluster (r= 0.61 and p =0.002). A similar correlation of the connectivity of the Left Thalamus to left Putamen, Pallidum, Caudate, Thalamus; Hippocampus and Insular Cluster [r= 0.55, p = 0.007] was noted (Figure 4B).

No statistically significant correlations of the age of onset of illness, the time interval between imaging, last seizures, age and gender with functional connectivity and network topology changes were noted.

## DISCUSSION

Febrile seizures are classified as generalised onset motor seizures in the revised ILAE classification of epilepsy^21^. The FEBSTAT study remarked that children presenting with Febrile Status Epilepticus are at risk for acute hippocampal injury in the background of structural imaging abnormalities ^22^. In this study, we demonstrate, that children with CFS have altered connectivity in the bilateral temporal lobes and thalami when compared to the SFS group. We also note increased segregation and decreased integration in several subcortical structures and the right frontal lobe in the CFS group. The increased connectivity of temporal lobes and decreased integration of the subcortical structures correlated with the number of recurrent febrile seizures, and the duration of the longest seizures^23,24^. While the causal nature and relevance of structural imaging findings are being contested ^25,26^, the findings of the current study add to the understanding of rs-fMRI-derived functional connectivity alterations in new-onset FS. We surmise that these changes may be the imaging translation of experimental evidence of alterations of functional neuronal and network properties (increased excitability) in early life epileptogenesis ^27,28^ without structural alterations such as neuronal death, genesis or altered branching ^29^. Increased excitability can in part explain reduced FC within and across circuits exhibiting this trait. With intact neurovascular coupling neurons with a differential higher likelihood of firing would lead to less coherent “ensemble type firing”/ local field potentials. Such less coherent (viz, noisier) neural activity can culminate in reduced RSFC correlation strength within and across resting state networks.

The functional connectivity imprint of febrile seizures reported in the current study could represent either the cause or effect of febrile seizures. Hyperconnectivity, a probable marker of increased neural resource use, is a well-documented ubiquitous response in many neurological disruptions ^30^. Notwithstanding its implications in terms of processing speed, cognitive fatigue and resource use ^31^, increased connectivity has nevertheless been noted as a pervasive reaction to many neurologic disturbances. Evidence suggests that this change in connectivity stems from increased spiking output from neuronal ensembles^32^. In the pathobiology of complex febrile seizures, the observed hyperconnectivity may have a role that is not necessarily compensatory. Enhanced EEG connectivity in febrile seizures has been incriminated as a seizure-prone state by Birca et al^33^. Mossy fibre plasticity and enhanced hippocampal excitability, without neither hippocampal cell loss nor altered neurogenesis, have been reported in animal models of prolonged febrile seizures^34^. Frequently described in the context of temporal lobe epilepsy, Kindling is a self-propagating reduction in the ability of the brain to limit seizures in which duration and behavioural involvement of induced seizures increase after repetitive induction of mesial temporal lobe seizures^35^. Sequential exaggerated mossy fibre invasion of the molecular and granule cell layers have been associated with the process of “kindling”. Murine models of hyperthermic seizures have documented evidence of the kindling-like phenomenon in epileptogenesis^36^. We propose that processes like these above-detailed mechanisms may operate in the genesis of hyperconnectivity in CFS as well the observed correlation with Febrile seizures’ recurrence and seizure duration.

Seizure circuitry in the context of temporal lobe epilepsy has been classified broadly as (a) the minimal-size initiating circuit(s) involving the trisynaptic circuit of the entorhinal cortex, the dentate gyrus(a “gatekeeper” resisting recruitment) and Ammon’s horn of the hippocampus and (b) the pathways of seizure spread by which additional brain circuits are recruited as the seizure continues and spreads^27^. In this context, it becomes difficult to ignore the similarity of the regions described in the experimental models with those obtained from the data-driven methodology used in the current study. The hyperconnectivity between allocortical [hippocampus, amygdala, accumbens] and neocortical regions [middle temporal gyrus] of bilateral temporal lobes is similar to the regions described in the seizure initiating circuit. There was also positive connectivity of bilateral thalami to multiple subcortical structures and cortical structures, probably indicating pathways of spread^37^.

Delving on the translational value from a clinical benefit perspective, the network topology of increased segregation and decreased integration found in our study indicates the simplified, regularised nature of these networks in these regions, reiterating the argument for disease-related network alterations (affecting long-range connection and forming self-reinforcing networks) in children with CFS. These observations are in tune with studies on subjects with generalised tonic-clonic epilepsy (affliction of bilateral temporal poles and thalamus)^38^ and benign epilepsy and centrotemporal spikes^39^. These structures have been fixated using other imaging approaches as well^40,41^. That the severity of network disturbances scales alongside variables such as duration of the seizure and number of recurrences only affirms the significance of these findings. Since the network alterations described herein in CFS is in children with “recent-onset” seizures, not confounded by other nuances of epilepsy (viz chronic drug intake), the association is likely “causative” and not an epiphenomenon/after-effect. Findings from another study in children with drug-naive recent-onset generalised epilepsy also supported thalamic atrophy as a cause and not the effect of epilepsy^42^. In addition, the nodal topology identified was a simplified pattern of the network, with increased segregation and decreased integration similar to the pattern observed in an immature brain^43^. The connectivity evidence from this study, first of its kind, makes a compelling case for reviewing the existing management pattern of patients with FS and advancing a body of research towards a well-differentiated treatment and follow-up algorithm.

This study is not without its limitations. It needs to be noted that imaging was performed in natural sleep due to ethical aspects of giving sedation in children undergoing imaging for febrile seizures. In this precarious situation, it was difficult to monitor the stages of sleep and hence the confounding effects of various stages of sleep on results cannot be eliminated. It is to be noted that the rs-fMRI pattern in sleeping infants closely resembles the adult sleep state rather than adult wakefulness ^25-12-2021^ 10:18:00 and few existing works of literature on network topology in sleep also reveal similar findings to the observations in the current study^44^. However, since both groups of children were sleeping and since the obtained results are in tune with the existing literature in other generalised motor epilepsies, it is more likely that the results are a genuine reflection of epilepsy than natural sleep. This study was not armed with a control group. For reasons similar to those cited above, the lack of a control group might also seem to be a limitation, but there is evidence for advantages of disease matched controls in evaluating heterogeneous diseases like epilepsy^18^. The results of our study are based on the group-level analysis between the two groups and might not be relevant to an individual patient. The sample size achieved was small. We lost significant data of nine children due to uncorrectable head motion due to snoring and hence the study sample size is small and generalizability to a larger population becomes difficult. The sampling time of rsfMRI data of 7 minutes is less than ideal to achieve stable RSFC data. Larger samples with longitudinal observations and the clinical outcome would add further evidence to the above observations.

## CONCLUSION

Children with recently diagnosed complex febrile seizures reveal altered connectivity having an immature simplified pattern in several regions including temporal lobes and thalami proportional to the frequency, and duration of the seizure. This evidence is in tune with experimental evidence in febrile seizures and the network topology in other generalised motor epilepsies, and hence more likely represent the cause of seizures. Regardless of the causal/consequential nature, such observations on altered connectivity demonstrate the imprint of these disease-defining variables of febrile seizures on the developing brain.

## MATERIALS AND METHODS

The prospective study was conducted at a tertiary care referral centre for neurologic disorders in children with recent-onset febrile seizures. Written informed consent was obtained from the caregiver of each participant, and the study was approved by the NIMHANS Human Ethics Committee-Basic and Neurosciences Division to be performed in children without using sedation. All the method were perfumed in accordance with the relevant guidelines and regulations. Hence all children underwent imaging while they were naturally sleeping inside the MRI gantry. This was associated with increased scan time, due to multiple pauses and restarts and resultant loss of data in nine subjects.

### Study population

Thirty-three patients with febrile seizures were recruited - [Simple FS - 19; Complex FS - 14]. Confirmation of the diagnosis of febrile seizures was based on clinical parameters and biochemical investigations according to the American Academy of Pediatrics guidelines^45^. After the exclusion of subjects with uncorrectable head motion, 24 subjects were included in the final analysis [Simple FS - 11 and Complex FS - 13]. The Male: Female ratio was 10:3 and 8:3 in the CFS and SFS groups respectively(p=1.0). A positive family history of seizures was present in four and three patients in the CFS and SFS groups respectively. Time interval from the last seizure to the MR imaging for the CFS and SFS groups were 12 days [IQR= 8.5 - 13.5] and 10 days [IQR = 9-30] respectively. Among the clinical variables analysed viz mean age at onset, the number of recurrent febrile seizures, the maximum duration of febrile seizures, duration of the disease, none showed significant between-group differences.

The demographic and clinical features of the patients are provided in Table 1

### Image acquisition

MRI was performed in a 3 Tesla scanner (SKYRA, Siemens, Erlangen, Germany). The child was allowed to sleep in the gantry room in the hands of the parent after ensuring that they both were metal-free after the room lights were dimmed. Once the child was asleep in the MR environment, a trained technologist transferred the child on the table and positioned with minimum disturbance to the sleeping child wrapped in a blanket. If at any point the child woke up, the entire cycle was repeated. The head was well supported with soft pads. 32 Channel head coil was used. The resting-state fMRI acquisition (rs-fMRI) using blood oxygen level-dependent (BOLD) contrast were as follows: 200 volumes, repetition time 2030ms, 40 slices, 3 mm slice thickness, FOV 195×195 mm, matrix 64×64, refocusing pulse 90°, voxel size-3 × 3 × 3mm The total time of acquisition for rs-fMRI was 6 minutes 52 seconds. Anatomic images were acquired by using a 3D T1-weighted MPRAGE sequence in 192 sagittal sections with a TR of 1900ms, TE of 2.5ms, a TI of 900ms, a FOV of 256 × 256 and a section thickness of 1mm. Oblique Coronal T2 Fast Spin Echo (FSE) planned perpendicular to the hippocampus was also done to rule out other structural abnormalities. Structural imaging revealed that one patient with complex febrile seizure had bulky and T2 hyperintense right hippocampus. The rest of the imaging studies did not reveal any abnormality.

### Image analysis

#### Pre-processing

The MRI data were pre-processed using MELODIC (Multivariate Exploratory Linear Optimized Decomposition into Independent Components) version 3.14, which is part of FMRIB’s Software Library (FSL, http://fsl.fmrib.ox.ac.uk/fsl). The pre-processing steps include: Discarding the first five functional images, removal of non-brain tissue, motion correction using MCFLIRT, intensity normalization, temporal band-pass filtering, spatial smoothing was applied using a 5mm FWHM Gaussian kernel, rigid-body registration was performed. The functional and structural data were co-registered to the 2year paediatric MNI template space (developed University of North Carolina [UNC]) using FLIRT (12 DOF) ^46^. Finally, single-session ICA with automatic dimensionality estimation was performed to identify noise components for each subject. Each ICA component was evaluated based on the spatial map, the time series, and the temporal power spectrum ^47^. Once the noisy ICA was marked by manual hand labelling, FIX was applied with default parameters to remove noisy components to obtain clean functional data. The head movement parameters (three translational and three rotational) were regressed out and 0.01 to 0.09Hz band pass filter was used before post processing.

#### Anatomic Parcellation

The fMRI data were segmented into 90 anatomic ROIs based on a University of North Carolina [UNC] paediatric [two years] atlas for whole-brain regions by using the anatomically labelled template reported by Shi et al, 2011^48^.

#### Functional Connectivity Analysis

A seed-to-voxel–based functional connectivity analysis was performed by computing the temporal correlation between the blood oxygen level-dependent signals to create a correlation matrix showing connectivity from a seed region to all other voxels in the brain by using the functional connectivity toolbox (CONN, version 17) implemented in SPM8 (http://www.nitrc.org/projects/conn)^49^. Meanwhile, source reduction of WM and CSF-related physiologic noises was carried out before connectivity estimation, by using the CompCor algorithm^50^. Bivariate correlations were analysed to reflect connections between the seed region to the rest of the brain voxels. Then Fisher’s r-to-z transformation was used on the connectivity matrix. This was followed by a general linear model that was designed to determine those statistically significant BOLD signal correlations between the mean time series from each seed ROI and that of every other brain voxel, at the individual subjects’ level (first-level analysis) ^51,52^.

Second-level random-effects analysis was used to create within-group statistical parameter maps for each network and to examine connectivity differences between groups. The group mean effects were estimated for both groups and seed to target connectivity was calculated using 2nd level co-variate analysis using CONN ^49^. Finally, Pearson linear correlation was performed between clinical variables [number of recurrent febrile seizures, the longest duration of febrile seizure, duration of disease, age of onset and time interval between last seizures and MRI] using the effect size of statistically significant seeds to target connectivity. The correlation coefficient, r (rho) and statistical significance, p values were calculated for each of these connectivity using MATLAB. Positive correlations were designated with a plus (“+”) sign and negative correlations with a minus (“-“) sign.

### Graph Theory Analysis

The graph-theoretical metrics were computed from the connectivity matrix. To this end, we extracted the BOLD time series for the parcellated 90 brain regions of interest (ROI). We followed this with an ROI-ROI rs-fMRI time-series correlation (Pearson correlation coefficients; individual subject) with a resultant 90*90 (n ROIs=90) connectivity matrix that was constructed. The functional connectivity brain-networks were defined based on 90*90 weighted undirected networks specified by G (N, E), where G is a non-zero subset with nodes (N=ROIs) and edges (E=inter-nodal correlation coefficients, Fisher’s Z values) to serve as a measure of functional connectivity between these nodes. We then computed the following graph theory measures: brain network segregation, small-worldness, network integration, and efficiency using the Brain connectivity toolbox (http://www.brain-connectivity-toolbox.net)^53,54^. Sparsity-based thresholding was employed to achieve a fixed, desired connection density to enable inter-group comparison.

### Brain Network Segregation

Network segregation was estimated from the Clustering Coefficient, which measures the strength of localised interconnectivity of a network. For a typical node (i)the absolute clustering coefficient (C_i_) is the ratio of the number of its existing connections (E) and all feasible connections of graph G. C is the average measure of all nodes.

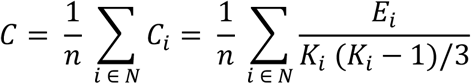

Where N is the total network nodes, E_i_ is the count of existent connections of node i, K_i_ being the node’s degree. Normalized whole-brain clustering coefficient (γ) was calculated from the ratio of absolute clustering coefficient of the network to random clustering coefficient (C_Rand_).

### Brain Network Integration

Participation coefficient measures the breadth or diversity of between-module connections of the individual nodes. Generally, nodes with a higher participation coefficient have a higher strength of connections to multiple modules. Participation coefficient (‘PC_i_’) of a node(i) is the ratio of ‘number of edges of node (i) to nodes in a module (s)’, and the degree of node ‘i’. Absolute participation coefficient may then be defined (PC_abs_) as:

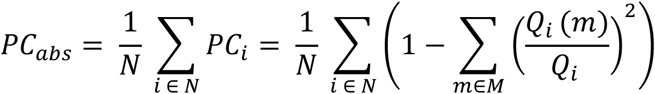

Wherein, ‘m’ is a module among a set of modules ‘M’, and ‘Qi (m)’ is the denoted by degree between node ‘ i’ to all other nodes in module ‘m’.

### Small-worldness

It is computed from the ratio between network segregation (γ) to that of network integration (λ). A network would be described as “small-world” if (i) γ > 1, and (ii) λ ≈ 1. Summarising into a simple measurement, σ = γ/λ > 1 for those networks that have a “small-world” organization ^55^.

### Whole Brain Efficiency

The brain network integration was measured using global efficiency, a measure of efficacy information exchange over the network^56^. It may be defined as an average of inverse shortest path length characterising that network ^57^. The higher the global efficiency of a network, the greater is the topological integration.

Finally, the brain regions that showed significant differences for on graph theoretical measures of segregation and integration were subsequently correlated with clinical measures. The brain regions that showed significant differences were rendered in a glass brain view using BrainNet Viewer (http://www.nitrc.org/projects/bnv/) ^58^.

### Statistical Analysis

Between groups (CFS and SFS) differences for the seed-to-voxel–based functional connectivity, seed to target connectivity was considered statistically significant with false discovery rate (FDR) corrected (p <0.05) at a clustered level for brain regions (N=90) using CONN^49^. Following this, we used a cluster size threshold of 20-voxels (i.e., if the connectivity cluster showed less than 20-voxels it was excluded).

For graph measure between groups (CFS and SFS) differences were assessed using a two-tailed two-sample t-test. The whole-brain graph measures statistical analysis was done at each Sparsity threshold with FDR correction for no of Sparsity (N=17). Brain regional segregation and integration differences were assessed for 90 brain regions using FDR correction with p<0.05 for no brain regions (N=90).

For clinical measures correlation with seed-to-voxel–based functional connectivity difference, we used FDR correction with p<0.05 for the total number of significant connectivity clusters for all the target ROIs (N=11). In the case of graph theory, we used FDR correction with p<0.05 for the total number of the brain regions that showed differences in regional integration or segregation (N=10).

## Acknowledgements

We thank the children with febrile seizures and their parents for being part of the study. We also thank the staff and students of the department of Neuroimaging and Interventional Radiology for their support during data acquisition.

## Author contributions

RDB, SS, BPS designed research. JS, AKG, KR, RCM, MLK recruited patients. UVA, KK, RP acquired and analysed the data. UVA, KK, RP and RDB wrote the manuscript. All authors contributed to result interpretation and editing of the manuscript.

## REFERENCES

1. Steering Committee on Quality Improvement and Management, Subcommittee on Febrile Seizures. Febrile Seizures: Clinical Practice Guideline for the Long-term Management of the Child With Simple Febrile Seizures. PEDIATRICS 121, 1281–1286 (2008).

2. Offringa, M. & Moyer, V. A. Evidence based paediatrics: Evidence based management of seizures associated with fever. BMJ 323, 1111–1114 (2001).

3. Stafstrom, C. E. Chapter 1 - The Incidence and Prevalence of Febrile Seizures. in Febrile Seizures (eds. Baram, T. Z. & Shinnar, S.) 1–25 (Academic Press, 2002). doi:10.1016/B978-012078141-6/50003-2.

4. Baulac, S. et al. Fever, genes, and epilepsy. Lancet Neurol. 3, 421–430 (2004).

5. Nakayama, J. Progress in searching for the febrile seizure susceptibility genes. Brain Dev. 31, 359–365 (2009).

6. Waruiru, C. Febrile seizures: an update. Arch. Dis. Child. 89, 751–756 (2004).

7. Berg, A. T. & Shinnar, S. Complex febrile seizures. Epilepsia 37, 126–133 (1996).

8. Jensen, F. E. & Sanchez, R. M. Chapter 11 - Why Does the Developing Brain Demonstrate Heightened Susceptibility to Febrile and Other Provoked Seizures? in Febrile Seizures (eds. Baram, T. Z. & Shinnar, S.) 153–168 (Academic Press, 2002). doi:10.1016/B978-012078141-6/50013-5.

9. Cendes, F. & Andermann, F. Early childhood prolonged febrile convulsions, atrophy and sclerosis of mesial structures, and temporal lobe epilepsy: 6.

10. Shinnar, S. & Glauser, T. A. Febrile seizures. J. Child Neurol. 17, S44–S52 (2002).

11. Theodore, W. H. Do Febrile Seizures Cause Mesial Temporal Sclerosis? Epilepsy Curr. 3, 121–122 (2003).

12. Pavlidou, E., Hagel, C. & Panteliadis, C. Febrile seizures: recent developments and unanswered questions. Childs Nerv. Syst. 29, 2011–2017 (2013).

13. Characteristics of medial temporal lobe epilepsy: I. Results of history and physical examination - French - 1993 - Annals of Neurology - Wiley Online Library. https://onlinelibrary.wiley.com/doi/abs/10.1002/ana.410340604.

14. Tsai, M.-L., Hung, K.-L., Tsan, Y.-Y. & Tung, W. T.-H. Long-term neurocognitive outcome and auditory event-related potentials after complex febrile seizures in children. Epilepsy Behav. 47, 55–60 (2015).

15. Dube, C. et al. Prolonged febrile seizures in the immature rat model enhance hippocampal excitability long term. Ann. Neurol. 47, 336–344 (2000).

16. Dubé, C. et al. Temporal lobe epilepsy after experimental prolonged febrile seizures: prospective analysis. Brain 129, 911–922 (2006).

17. Theodore, W. H. et al. Hippocampal atrophy, epilepsy duration, and febrile seizures in patients with partial seizures. Neurology 52, 132–132 (1999).

18. Bharath, R. D. et al. Seizure Frequency Can Alter Brain Connectivity: Evidence from Resting-State fMRI. AJNR Am. J. Neuroradiol. 36, 1890–1898 (2015).

19. Bharath, R. D. et al. Reduced small world brain connectivity in probands with a family history of epilepsy. Eur. J. Neurol. 23, 1729–1737 (2016).

20. Bernhardt, B. C., Bonilha, L. & Gross, D. W. Network analysis for a network disorder: The emerging role of graph theory in the study of epilepsy. Epilepsy Behav. 50, 162–170 (2015).

21. Fisher, R. S. et al. Operational classification of seizure types by the International League Against Epilepsy: Position Paper of the ILAE Commission for Classification and Terminology. Epilepsia 58, 522–530 (2017).

22. Shinnar, S. et al. MRI abnormalities following febrile status epilepticus in children: The FEBSTAT study. Neurology 79, 871–877 (2012).

23. Silverstein, A. M. & Alexander, J. A. Acute postictal cerebral imaging. Am. J. Neuroradiol. 19, 1485–1488 (1998).

24. Szabo, K. et al. Diffusion-weighted and perfusion MRI demonstrates parenchymal changes in complex partial status epilepticus. Brain 128, 1369–1376 (2005).

25. Grillo, E. & Ronaldo J. M. da Silva, F. MRI peri-ictal abnormalities in febrile status epilepticus. Cause or consequence? (2020).

26. Berg, M. J. & Abou-Khalil, B. Childhood febrile status epilepticus: Chicken or egg? Does it matter? Neurology 79, 840–841 (2012).

27. Bertram, E. The relevance of kindling for human epilepsy. Epilepsia 48 Suppl 2, 65–74 (2007).

28. Lothman, E. W., Bertram, E. H. & Stringer, J. L. Functional anatomy of hippocampal seizures. Prog. Neurobiol. 37, 1–82 (1991).

29. McClelland, S., Dubé, C. M., Yang, J. & Baram, T. Z. Epileptogenesis after prolonged febrile seizures: Mechanisms, biomarkers and therapeutic opportunities. Neurosci. Lett. 497, 155–162 (2011).

30. Hillary, F. G. et al. Hyperconnectivity is a fundamental response to neurological disruption. Neuropsychology 29, 59–75 (2015).

31. Nakamura, T., Hillary, F. G. & Biswal, B. B. Resting network plasticity following brain injury. PloS One 4, e8220 (2009).

32. Chawla, D., Lumer, E. D. & Friston, K. J. Relating macroscopic measures of brain activity to fast, dynamic neuronal interactions. Neural Comput. 12, 2805–2821 (2000).

33. Birca, A. et al. Enhanced EEG connectivity in children with febrile seizures. Epilepsy Res. 110, 32–38 (2015).

34. Bender, R. A., Dubé, C., Gonzalez-Vega, R., Mina, E. W. & Baram, T. Z. Mossy fiber plasticity and enhanced hippocampal excitability, without hippocampal cell loss or altered neurogenesis, in an animal model of prolonged febrile seizures. Hippocampus 13, 399–412 (2003).

35. Goddard, G. V., McIntyre, D. C. & Leech, C. K. A permanent change in brain function resulting from daily electrical stimulation. Exp. Neurol. 25, 295–330 (1969).

36. Dayao Zhao, Xiru Wu, Yinquan Pei, & Qihua Zuo. Kindling phenomenon of hyperthermic seizures in the epilepsy-prone versus the epilepsy-resistant rat. Brain Res. 358, 390–393 (1985).

37. Danielson, N. B., Guo, J. N. & Blumenfeld, H. The default mode network and altered consciousness in epilepsy. Behav. Neurol. 24, 55–65 (2011).

38. Li, Y., Chen, Q. & Huang, W. Disrupted topological properties of functional networks in epileptic children with generalized tonic-clonic seizures. Brain Behav. 10, e01890 (2020).

39. Ji, G.-J. et al. Decreased Network Efficiency in Benign Epilepsy with Centrotemporal Spikes. Radiology 283, 186–194 (2017).

40. Huang, W. et al. Gray-matter volume reduction in the thalamus and frontal lobe in epileptic patients with generalized tonic-clonic seizures. J. Neuroradiol. J. Neuroradiol. 38, 298–303 (2011).

41. Wang, Y., Goodfellow, M., Taylor, P. N. & Baier, G. Dynamic Mechanisms of Neocortical Focal Seizure Onset. PLOS Comput. Biol. 10, e1003787 (2014).

42. Perani, S. et al. Thalamic volume reduction in drug-naive patients with new-onset genetic generalized epilepsy. Epilepsia 59, 226–234 (2018).

43. Fair, D. A. et al. The maturing architecture of the brain’s default network. Proc. Natl. Acad. Sci. 105, 4028–4032 (2008).

44. Lv, J., Liu, D., Ma, J., Wang, X. & Zhang, J. Graph Theoretical Analysis of BOLD Functional Connectivity during Human Sleep without EEG Monitoring. PLOS ONE 10, e0137297 (2015).

45. Management, S. C. on Q. I. and & Seizures, S. on F. Febrile Seizures: Clinical Practice Guideline for the Long-term Management of the Child With Simple Febrile Seizures. Pediatrics 121, 1281–1286 (2008).

46. Jenkinson, M., Bannister, P., Brady, M. & Smith, S. Improved optimization for the robust and accurate linear registration and motion correction of brain images. NeuroImage 17, 825–841 (2002).

47. Beckmann, C. F., DeLuca, M., Devlin, J. T. & Smith, S. M. Investigations into resting-state connectivity using independent component analysis. Philos. Trans. R. Soc. Lond. B. Biol. Sci. 360, 1001–1013 (2005).

48. Shi, F. et al. Infant Brain Atlases from Neonates to 1- and 2-Year-Olds. PLoS ONE 6, e18746 (2011).

49. Whitfield-Gabrieli, S. & Nieto-Castanon, A. Conn: A Functional Connectivity Toolbox for Correlated and Anticorrelated Brain Networks. Brain Connect. 2, 125–141 (2012).

50. Behzadi, Y., Restom, K., Liau, J. & Liu, T. T. A Component Based Noise Correction Method (CompCor) for BOLD and Perfusion Based fMRI. NeuroImage 37, 90–101 (2007).

51. Whitfield-Gabrieli, S. & Ford, J. M. Default mode network activity and connectivity in psychopathology. Annu. Rev. Clin. Psychol. 8, 49–76 (2012).

52. Lindquist, M. A., Loh, J. M., Atlas, L. Y. & Wager, T. D. Modeling the Hemodynamic Response Function in fMRI: Efficiency, Bias and Mis-modeling. Neuroimage 45, S187–S198 (2009).

53. Rubinov, M. & Sporns, O. Complex network measures of brain connectivity: Uses and interpretations. NeuroImage 52, 1059–1069 (2010).

54. Zalesky, A., Fornito, A. & Bullmore, E. T. Network-based statistic: Identifying differences in brain networks. NeuroImage 53, 1197–1207 (2010).

55. Bassett, D. S. & Bullmore, E. D. Small-world brain networks. The neuroscientist 12, 512–523 (2006).

56. Iturria-Medina, Y., Sotero, R. C., Canales-Rodríguez, E. J., Alemán-Gómez, Y. & Melie-García, L. Studying the human brain anatomical network via diffusion-weighted MRI and Graph Theory. NeuroImage 40, 1064–1076 (2008).

57. Latora, V. & Marchiori, M. Efficient Behavior of Small-World Networks. Phys. Rev. Lett. 87, 198701 (2001).

58. Xia, M., Wang, J., & He, Y. BrainNet Viewer: a network visualization tool for human brain connectomics. PloS one 8, e68910 (2013).

